# *In silico* assessment of the association of pathogenicity and metal-resistance potential of *Fusarium* spp

**DOI:** 10.1101/2022.10.12.511937

**Authors:** Gerald Amiel M. Ballena, Michael C. Velarde, Leilani S. Dacones

## Abstract

Genus *Fusarium* includes ubiquitous species complexes and are often resistant to multiple stressors. Early epidemiological evidence show that metal resistance genes (MRGs) influence the spread of antibiotic resistance genes (ARGs) in microbial communities. More recent evidence points out that this correlation is due to the physical linkage of these genes. Given the utmost importance of *Fusarium* pathogenicity to agriculture, and the ever-increasing rise in metal or metalloid displacement in the environment – this paper aims to pioneer the investigation of whether pathogenicity determinants also correlate well with MRGs. To provide probable patterns of horizontal gene transfer or incomplete lineage sorting, a species tree was initially defined. *Fusarium* is unanimously monophyletic from our phylogenetic analysis of 19 concatenated loci. However, saturation analysis show that most of sequences beyond the Terminal Fusarium Clade (TFC) are saturated and are likely to show erroneous phylogenetic relationships. Further analysis of tree topologies also show discordance among loci. Metal-resistance proteins (MRPs) and pathogenicity-related proteins (PRPs) were present in almost all the genomes tested. Remarkably, correlation between MRPs and PRPs among *Fusarium* is positive and statistically significant. Moreover, there the some of the MRPs and PRPs co-occur significantly more than chance alone. Overall, this suggests that there is a possibility that pathogenicity and metal tolerance proteins among *Fusarium* may co-occur.

## Introduction

*Fusarium* refers to a broad monophyletic group of ubiquitous Ascomycota fungi which encompass medically and agriculturally important strains (Geiser, et al., 2021; Crous, et al., 2021; O’Donnell, et al., 2020). Given the importance of *Fusarium* to agriculture, a robust classification system is needed for us to differentiate pathogenic from non-pathogenic cryptic forms. Traditionally, phylogenetic classification was done using morphological, biochemical, and physiological characteristics. While more recent classifications schemes also consider genetic or genomic characters (e.g., Crous, et al., 2021). The consensus is that *Fusarium* consists of a well-supported Terminal Fusarium Clade (TFC) and an ill-delineated Basal Fusarium Clade (BFC) (Geiser, et al., 2021; Crous, et al., 2021). Both studies made use of a 19-exon tree for phylogenomic resolution because sampling a larger number of genes can help resolve conflict among individual genes.

However, mounting evidence show that pathogenicity of *Fusarium* spp. does not correlate well with the evolution of housekeeping genes (Taylor, et al., 2016; Williams, et al., 2016) – i.e., they appear polyphyletic with respect to species trees. Polyphyly in phylogenetics can arise from different evolutionary events such as incomplete lineage sorting (ILS), introgression, hybridization, selection pressure, and Horizontal Gene Transfer (HGT).

To infect plants, plant-pathogens suppress or evade plant immune responses using a set of proteins called effectors (Pieterse, et al., 2009; De Wit, et al., 2009; Stergiopoulos & de Wit, 2009). Most effectors are in accessory chromosomes (ACs) (e.g., van der Does, et al., 2016). ACs also known as dispensable chromosomes do not contain genes that are essential for basal functions. These chromosomes often have more polymorphisms, repetitive sequences (Inami, et al., 2012; Achari, et al., 2021; Peng, et al., 2019), duplication events (Armitage, et al., 2018), asymmetric distribution among lineages (Jurgenson, et al., 2002), and pathogenic phenotypes (Fones and Gurr, 2017; Ma, et al., 2010; Peng, et al., 2019) – which make ACs can differ considerably terms genetically and epigenetically with the core genome (Yang, et al., 2020). These more chaotic regions of the genome allow for faster adaptation and a higher risk of emergence of pathogenicity (Schmidt, et al., 2013). Determining the evolution of PRPs is essential for understanding how phytopathogens spread. Thus, efforts to characterize and catalogue the repertoire of pathogenicity-related proteins (PRPs) are underway (e.g., PHI-base).

With the advent of next generation sequencing, more and more sequences are included in public databases – faster than they are processed. This allowed us to investigate pathogen emergence at an unprecedented rate. With this technology we now know that HGT, hybridization, and genomic plasticity all influence pathogen emergence (Thynne, et al., 2015). All of which lead to a polyphyletic distribution with respect to the species tree. Moreover, comparative genomics reveal that greater conservation exists among different *formae speciales* (Williams, et al., 2016). Which means that we can survey potential pathogenic phenotypes by determining the allelic repertoire of *Fusarium* species. Studying these pathogenic genotypes will allow us to determine the risk of an impending epidemic.

Aside from pathogenicity, a fast-evolving adaptive genome likely allowed *Fusarium* to be well-adapted in various stressful environments. With the increasing demand for metal and metalloids for technology, we see an increase in mining pollution of pristine environments. Recent papers have shown that *Fusarium* is able to colonize high-metal environments (e.g., Velmurugan, et al., 2010; Bonilla et al., 2021). Due to their agricultural importance, many well-documented sequences of fusaria can be found in online repositories. While many of these sequencing efforts exists, none so far have been used to determine the correlation or co-occurrence of metal resistance and pathogenicity.

Thus, the main goal of this study was to investigate whether a co-existence or cooccurrence between MRPs and PRPs exist within fusaria exist using previously published sequences. We produced a genus-level phylogeny of the TFC using various phylogenomic approaches to obtain a reference species trees – with which we can compare the trajectories of MRP (metal resistance proteins) and PRP (pathogenicity-related proteins) genes. Making use of sequences from public databases also allowed us to investigate the monophyly of *Fusarium* using the 19 exons proposed by Geiser, et al., (2021). We then compared our results with that of Geiser et al., (2021) and Crous, et al., (2021) with regards to the TFC.

## Materials and Methods

### Sequence data and Homology detection

All sequences from the manually curated databases PHI-base (Urban, et al., 2019) and BacMet (Pal, et al., 2013), were used as query sequences. A total of 102 representative assemblies from Hypocreales were used in this study, 95 of which were from Nectriaceae and from which 74 were from genus *Fusarium* (Table S1).

Phylogenetic trees were rooted to the members of Nectriaceae (*Stylonectria norvegica* [Snor; Crous, et al., 2021]) or Hypocreales (*Stachybotrys chlorohalonata* [Schl; Lombard, et al., 2015; Montoya, et al., 2021], *Sarocladium implicatum* [Simp; Summerbell, et al., 2011]) that are known to be most distant from *Fusarium* because *Fusarium* is classified under family Nectriaceae and order Hypocreales. All *Fusarium* proteome and genome data were downloaded from NCBI Datasets (https://www.ncbi.nlm.nih.gov/datasets/genomes/).

All similarity-based searches were given a cut-off of E-value > 1e-15. OrthoFinder (Emms and Kelly, 2019) was used to determine orthogroups using proteome sources.

### Phylogenetic resolution

The nineteen exon (19) loci previously used by Geiser, et al. (2021) and Crous et al. (2021) in testing the circumscription of *Fusarium* were used to determine the species tree in this current study (Table S2). Homologous nucleotide sequences were taken from the annotated genome sequences using BLASTN (-task blastn option) from the ncbi-blast 2.13.0+ package. For multiple BLAST hits, the hit with the highest bit score was considered the homolog. Additionally, 3498 genes containing exactly one sequence per individual (strict definition) (single-copy orthogroups) taken from OrthoFinder (Emms and Kelly, 2019) were also used create single-loci and coalescent species trees. All single copy loci were then aligned with MAFFT 7.490 (Katoh and Standley, 2013). To increase alignment accuracy, all alignments were refined using L-INS-I option.

Sequences with unique gaps (> 100 bp total) were removed from multiple sequence alignments (MSAs) as partial sequences can bias tree inference more than missing loci (Hosner, et al., 2015). To increase the reliability of trees to represent evolutionary history, pairwise transition (Ti) and transversion (Tv) plots under the GTR model were generated using DAMBE 7 (Xia, 2018) to determine substitution saturation. Sequences with the highest pairwise Tv > Ti values were iteratively removed until Tv values no longer exceeded Ti values i.e., Tv – Ti < 0. After these filtering steps, gaps were removed and sequences were re-aligned using MAFFT 7.490 (Katoh and Standley, 2013) to form ‘relaxed’ MSAs. Representative species without any missing loci were then extracted and used to generate a separate ‘strict’ alignment. A summary of all sequences trimmed due to either 1) the presence of gaps or 2) substitution saturation is in the figure below. Sequences with the complete 19-locus set is also shown (Figure S1).

### Tree building

Multiple sequence alignments (MSA) of each locus were used for either concatenated or coalescence-based tree building methods. Single loci were concatenated using DAMBE 7 (Xia, 2018). The best substitution model was identified using ModelFinder (Kalyaanamoorthy, et al., 2017) for each individual locus. All substitution models were selected using Bayesian Information Criterion (BIC). Partitioning of concatenated alignments were defined by the full lengths of each locus (Table S2). Maximum likelihood trees of both single and concatenated loci were constructed using IQ-tree2 (Minh, et al., 2020). Individual loci were treated as individual partitions. Branch support was evaluated using 1000 UFBoot2 replicates (Hoang, et al., 2017). Bayesian inference (BI) was performed using MrBayes 3.2.7 (Ronquist & Huelsenbeck, 2003; Altekar et al., 2004) with four Markov chains (3 heated), sampled every 5000 generations. A 16-category gamma distribution was used to accommodate rate heterogeneity among sites. Individual loci were run for 10,000,000 generations. Convergence was assessed using RWTY (Warren, et al., 2017). Distancebased minimum evolution (ME) trees were built from FastME (Lefort, et al., 2015) which uses F84 as the DNA evolutionary model and LG for protein by default, and maximum parsimony (MP) were constructed from mpboot (Hoang, et al., 2018). MP and ME trees were resampled with 1000 standard bootstrap replicates. Additionally, trees generated from the STAG (Emms & Kelly, 2018) and STRIDE (Emms & Kelly, 2017) algorithms were used as part of OrthoFinder (Emms & Kelly, 2019). Individual loci were then summarized using ASTRAL (Rabiee, et al., 2019). Trees were rooted based on previous literature. A summary of all trees generated in this study is shown in Figure S2.

### Detection of MRPs and PRPs

To determine the repertoire of metal resistance proteins (MRPs) and pathogenicity related proteins (PRPs) among *Fusarium* species and strains, all PHI-base (Urban, et al., 2019) and BacMet (Pal, et al., 2013) sequences were retrieved for similarity searching. All protein sequences were queried using DIAMOND (Buchfink, et al., 2021) against a local concatenated database of all predicted protein sequences from the *Fusarium* assemblies (see Sequence Data and Homology Detection). To ensure that all pairwise hits are the most likely orthologous only the hits from *Fusarium* sequences with the highest bit score was kept for further analyses.

For pathogenicity-related proteins, hits were removed if they are part of the “core” proteome set. Given that the core genome is defined as the set of genes that are ‘indispensable’ – for this study, the core proteome set was defined as the set of all proteins that belong to an orthogroup that is present in all representatives of each species complex. In more detail, all orthogroups were defined using OrthoFinder (Emms & Kelly, 2019). Using this data, orthogroups present in all representatives of a species complex (i.e., FOSC, FFSC, FsambSC, FSSC, and FIESC) were defined as the core-proteome for that species complex. The core proteomes of all species complexes were then clustered to define the core proteome set. This conservative approach was done to ensure that all indispensable proteins that define each species complex are included. All homologous sequences for each gene at this step were then used to create binary (presence/absence) heatmaps. Homologous sequences were aligned using MAFFT v 7.490 (Katoh and Standley, 2013).

### Topology and exploration of trees

To characterize the heterogeneity of the 19 species-defining loci (Geiser, et al., 2021), all the trees generated from MrBayes (Ronquist & Huelsenbeck, 2003) were used to produce the of BI-consensus trees (burn-in 25% + 1; 15,000 total trees). This conservative-burn-in value was also used to ensure that the trees have already reached convergence as stated by RWTY (Warren, et al., 2017). To lessen computational burden, only every 75 trees were sampled post burn-in totaling 3800 trees (200 trees x 19 loci). We also characterized the heterogeneity of ML trees created from MAFFT-aligned (Katoh and Standley, 2013) single-copy orthologs defined by OrthoFinder (Emms and Kelly, 2019). Gene tree heterogeneity is visualized using either principal coordinates analysis (PCoA) via treespace package (Jombart, et al., 2017). Shimodaira-Hasegawa and the Approximately Unbiased (AU) topology tests (Shimodaira & Hasegawa, 1999; Shimodaira, 2002) were performed in IQ-tree2 (Minh, et al., 2020) to assess statistical significance of tree incongruence. The ML concatenated alignment is the approach mostly used in literature and was thus used as base alignment.

### Genetic co-occurrence analysis

To resolve the co-occurrence of PRPs and MRPs, all orthologs detected by DIAMOND (Buchfink, et al., 2021) were converted into binary (absence/presence) matrices. Some sequences in BacMet (Pal, et al., 2013) correspond to the same protein, species with multiple putative copies of protein are only considered to have a “present” value in this matrix. Both PRP and MRP matrices were clustered using the Unweighted Pair Group Method with Arithmetic Mean (UPGMA) using the hclust function in R (R Core Team, 2022). To explore pairwise differences in UPGMA and phylogenetic tree topologies, tanglegrams were generated using the phytools package (Revell, 2011).

To determine whether the observed co-occurrences between different PRPs and MRPs in the 34 representative species is higher than expected from random chance, package cooccur (Veech, 2013) was used on the binary presence/absence matrix of all PRPs and MRPs. Each pair that cooccurs at a frequency higher than expected from random chance (p < 0.01) were then visualized in a co-occurrence network. Networks were generated using ggraph (Pedersen, 2021), igraph (Csardi & Nepusz, 2006), and RColorBrewer (Neuwirth, 2022).

### Statistical analyses and data visualizations

Total number of PRPs and MRPs were counted for each of the 34 representative sequences. Based on the PRP and MRP counts, the Pearson correlation coefficient was calculated using R (R Core Team, 2022) and plotted using ‘ggpubr’ R package (Kassambara, 2020). Histograms were generated in R. Binary heatmaps, bubble plots, and bar graphs were created using ggplot2 (Wickham, 2016). Phylogenetic trees were visualized and combined using TreeGraph2 (Stöver & Müller, 2010). Maps were processed using package rgdal (Bivand, et al., 2022) and ggplot2 (Wickham, 2016).

## Results and Discussion

### Confirmation of Terminal Fusarium Clade Phylogeny

The genus *Fusarium* is well-defined monophyletic group at the Terminal Fusarium Clade (TFC) separate from other members of Nectriaceae as shown in Figure 1 below. Within this clade, *Fusarium oxysporum* species complex (FOSC) is sister to *Fusarium fujikuroi* while *Fusarium incarnatum-equiseti* species complex is sister to *F. sambucinum. Fusarium solani* species complex is sister to all other *Fusarium* species complexes. These groupings are consistent with previous phylogenomic inferences (Crous, et al., 2021; Geiser, et al., 2021). Ultimately, this result means that the 19-exon concatenated tree is robust in delineating groups in TFC using our dataset (Table S1).

**Figure 1.**
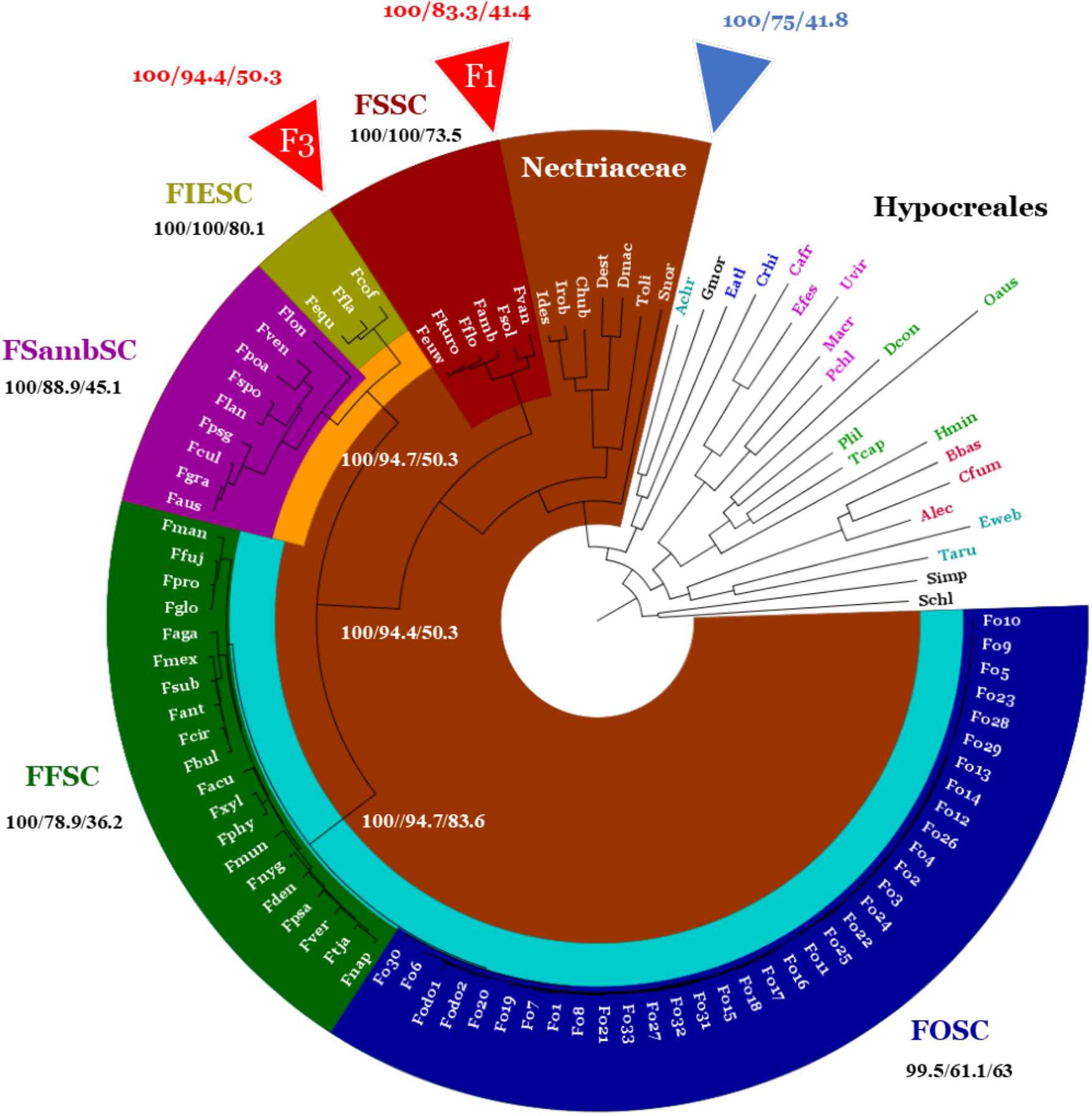
Complete concatenated ML tree. Colors represent different clades and subclades namely: blue – *Fusarium oxysporum* species complex, green – *Fusarium fujikuroi* species complex, pink – *Fusarium sambucinum* species complex, yellow – *Fusarium incarnatum*-*equiseti* species complex, red – Fusarium *solani* species complex and brown – *Nectriaceae*. Non-highlighted clades various species from Hypocreales. Light blue represents the FOSC and FFSC sister groups, while orange represent FsambSC and FIESC. Relaxed trees were rooted on *Stachybotrys chlorohalonata* (Schl; Lombard, et al., 2015; Montoya, et al., 2021) and *Sarocladium implicatum* (Simp; Summerbell, et al., 2011). White values show the supports for each sister-taxa relationship i.e., FFSC-FOSC, FsambSC-FIESC, ingroup Fusarium. Black support values indicate monophyletic support. Red values indicate the values for the F1 and F3 nodes, respectively. Blue values indicate support for the monophyly of *Nectriaceae*.

**Figure 2.**
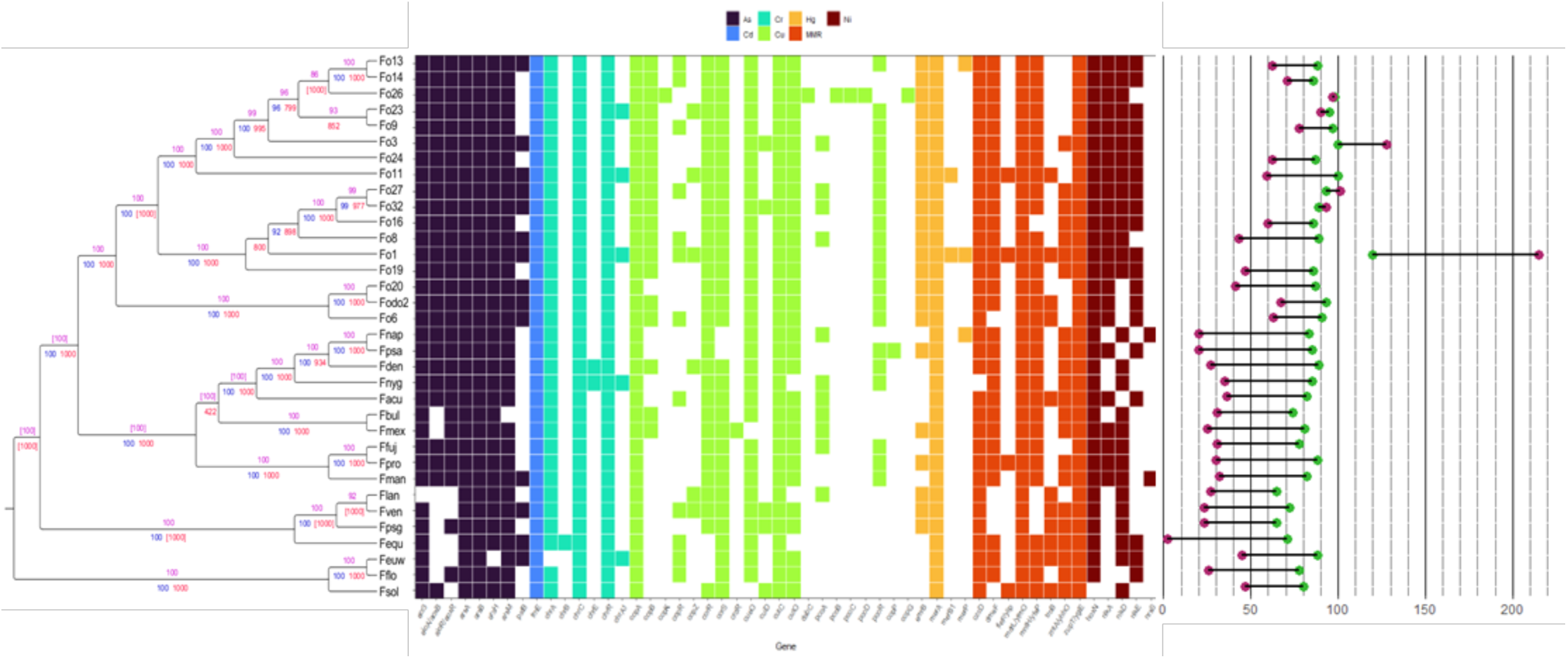
Presence or absence of selected metal-resistance proteins (MRPs) in various *Fusarium* strains. Phylogenetic tree used for this figure was the concatenated ML tree of the “strict” alignment. Values on the concatenated-ML tree represent support values supporting or conflicting with the clade assignment. Conflicting values are enclosed in square brackets ‘[]’. The support values include UFBoot values > 95 (blue), Bayesian posterior values (expressed as percentage; pink), and ME bootstraps (out of n=1000 bootstrap replicates, red). Presence of homologous sequences are denoted by colored tiles. Tiles are colored according to the resistance conferred by each MRP. The absence of a homolog is denoted by non-colored tiles. Tiles are colored according to the resistance conferred by each MRP. The absence of a homolog is denoted by non-colored tiles. The dumbbell plots on the right show the number MRPs (green) and PRPs (red) per respective species on the phylogenetic tree.

However, many of these sequences were saturated outside of the TFC (Figure S1) – which would have obscured the true phylogenetic distances due to long-branch attraction (Bergsten, 2005; Philippe, et al., 2009) or mislead relationships among groups (e.g., Breinholt & Kawahara, 2013). Moreover, the sequences themselves may have had gaps that might lead to erroneous inferences. To tackle this, we removed species-sequences that do not have any of the loci of the nineteen (19) exons described by Geiser, et al., (2021) detected by BLASTN. We also removed species that contained saturated loci to improve inferences (Duchêne et al., 2021). A presence/absence heatmap of the 19 exons can be seen in (Figure S1).

Notably, single- and multi-gene phylogenetic datasets can conflict with each other due to various sources of data error or conflicts/lack of phylogenetic signal (Philippe, et al., 2009). Current classification of *Fusarium* species is primarily based on concatenated alignment of several conserved nuclear loci – which, happen to correlate well with morphological and physiological data (Crous, et al., 2021). Concatenation assumes that all these loci are passed down as a supergene – which is likely because they are all important for immediate survival. To explore this, we used several tree building methods using the “strict” alignment to see whether other algorithms would differ in topology (Figure S2). Principal Coordinates Analysis (PCoA) and statistical topology tests of different trees generated from the 19-loci were showed that there were visual and statistically significant differences between tree topologies (Figure S3 and Table S3). This means that the 19 loci have differing phylogenetic signals and inferred evolutionary histories. Which ultimately implies that detection of polyphyly of PRPs and MRPs with respect to the species tree can differ depending on the loci and/or tree building methods used.

### *Fusarium* species have a diverse set of MRPs and PRPs

In contrast to the genes used in species delineation, effectors and metal-resistance determinants are not necessary for immediate survival – instead, they are required adaptation. To study the potential of the strains to resist metal and incur pathogenicity, proteomes of “strict” (see above) alignments were subjected to DIAMOND (Buchfink et al., 2021) many-to-many similarity searches. Results of the similarity searches were summarized using an absence/presence matrix as shown below (Figure 3 and Figure 4).

**Figure 3.**
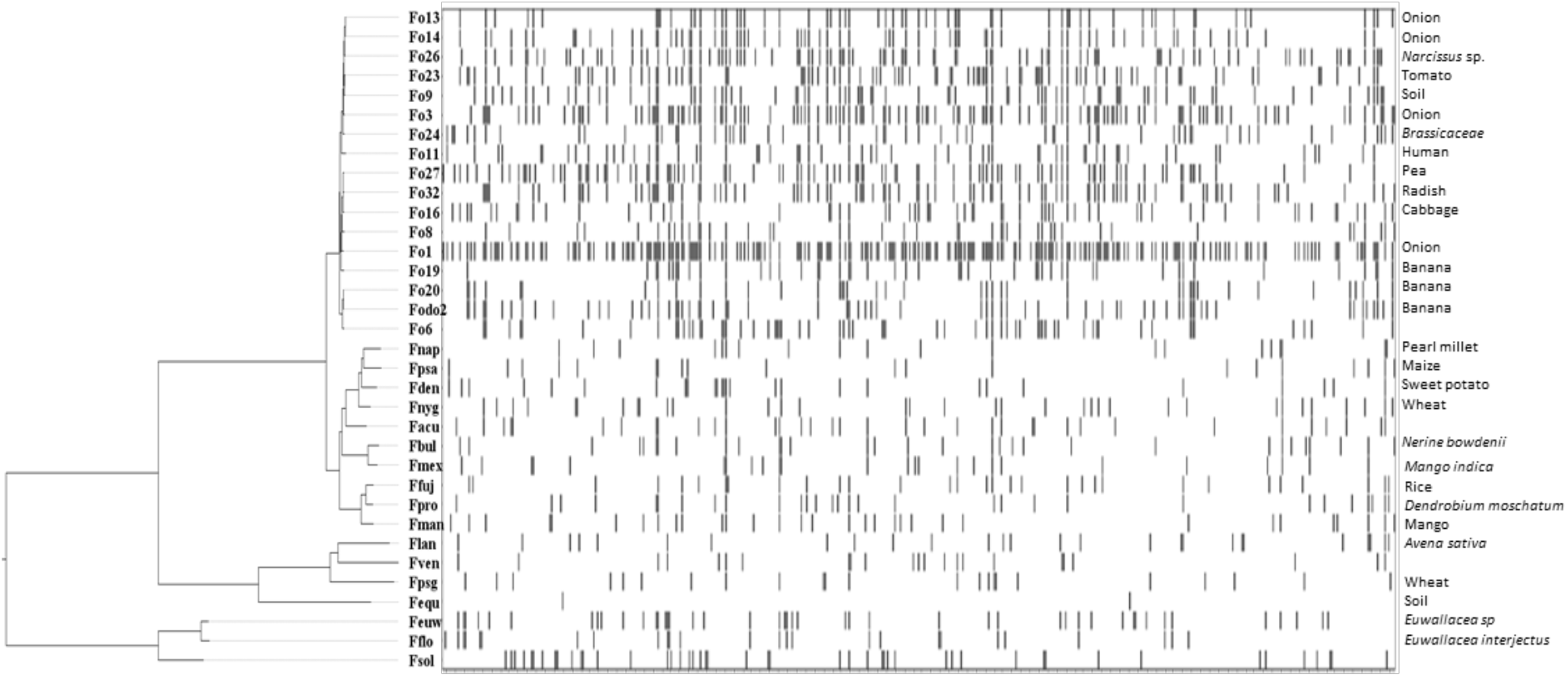
Presence or absence heatmap of pathogenicity-related proteins in Fusarium. Blackened tiles represent the presence of a homolog from each respective proteome indicated on the y-axis. Labels on the right indicate the putative hosts of each strain as indicated in their BioSample data.

**Figure 4.**
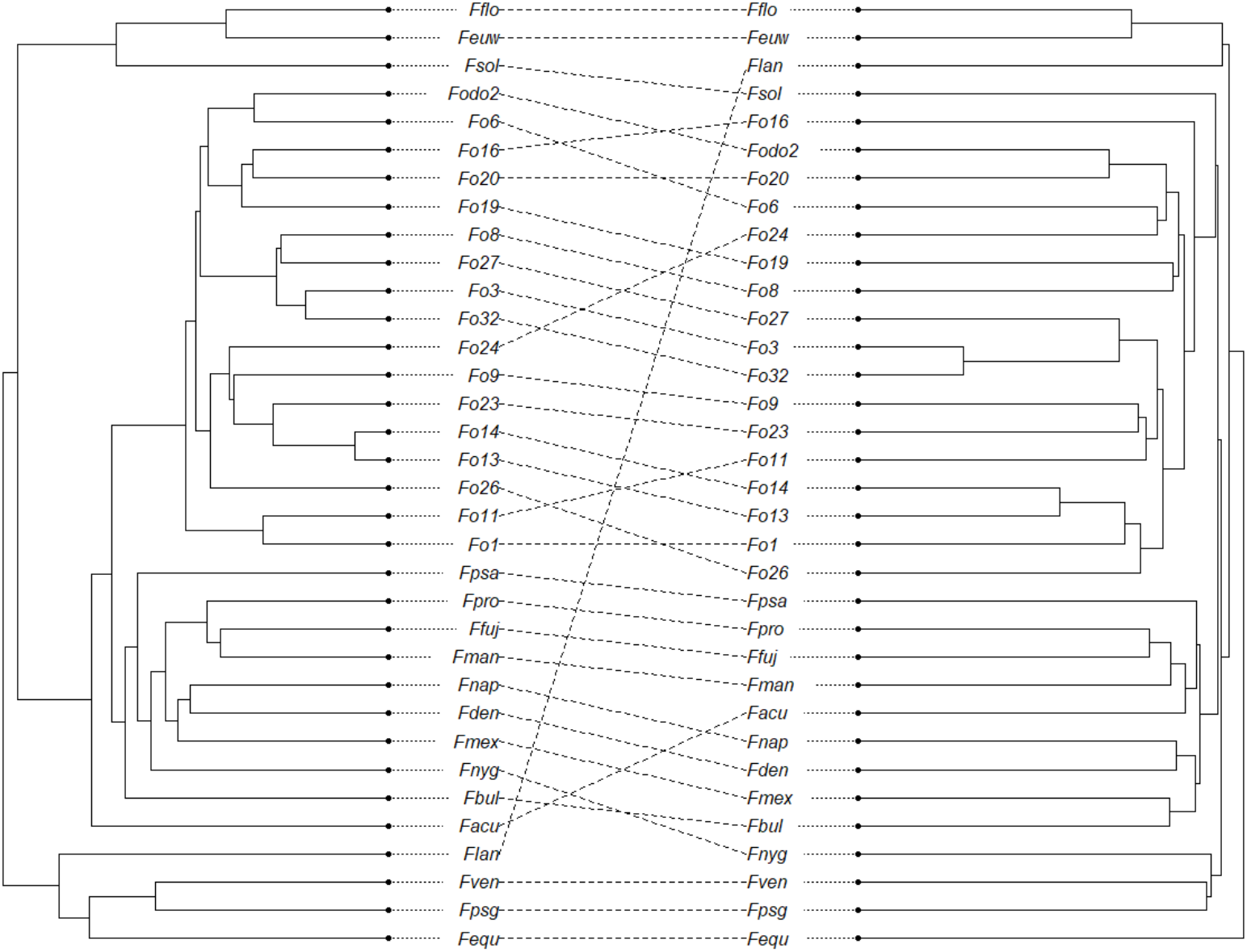
Tanglegram of rooted MRP (left) vs PRP (right) genotypes generated using phytools (Revell, 2011).

Immediately, we can observe that most of the *Fusarium* proteomes contain many hits with Arsenic-resistance and Copper-resistance proteins. This is generally what we would expect due to the widespread adaptation of life to Arsenic (Chen, et al., 2017; Danczak, et al., 2019) and Copper (Decaria, et al., 2011; Li, et al., 2015). From (Figure SX), we can also observe that, for most of the fusaria, there are more MRPs than PRPs. The relatively constant length of the dumbbell plots also hints a possible correlation between the number of MRPs and PRPs present in each strain.

Discarding the ‘core’ genome can be an effective strategy when looking for pathogenicity-related effectors (Williams, et al., 2016; van Dam, et al., 2016). Thus, to define a core set of proteins, we used OrthoFinder (Emms and Kelly, 2019) to look for orthogroups from the downloaded proteome (see Methods) set. A total of 508,768 out of 604,297 proteins were part of the ‘core’ proteome set – which accounts for about 84.19% of the whole proteome set. Figure 4 below shows the nonredundant DIAMOND hits of PRPs presented as a binary absence/presence matrix.

Figure 3 highlight the polyphyly of pathogenicity with respect to the concatenated-ML species tree. Notably, there are also noticeable differences between the ‘genotype’ of each strain – more so compared MRPs. Most notable are the placements of Flan, Facu, Fnyg, Fo11 and Fo26. In terms of phylogeny, both MRP and PRP UPGMA genotypes had discrepancies with the concatenated ML species tree (Figure S4 and S5). These results showcase discordance between the species tree and UPGMA genotypes. A tanglegram of between MRP and PRP UPGMA trees (Figure 4) suggest that there is also discordance between the metal-resistant and pathogenicity genotypes. Despite this, there is a significant positive correlation between the number of PRPs and MRPs (Figure S6).

### Co-occurrence between MRPs and PRPs

Figure 5 below is a network diagram showing the relationship between different clusters of proteins used in this study. Each connection in the diagram is plotted when the pairwise nodes cooccur more than random chance (Veech, 2012). This result highlighted the possibility that some of the major MRP groups do co-occur with themselves, and with other PRP groups. Further network analysis (Figure 6) below shows that there are indeed statistically significant cooccurrences between specific MRPs and PRPs. Considering that many of these species were sampled independently, this implies that either there is mixing among lineages or selection pressure to keep both types of proteins. More robust genotypic analysis would be able to explain the causes of phylogenetic and ‘genotypic’ discordance and the significant co-occurrence patterns between MRPs and PRPs.

**Figure 5.**
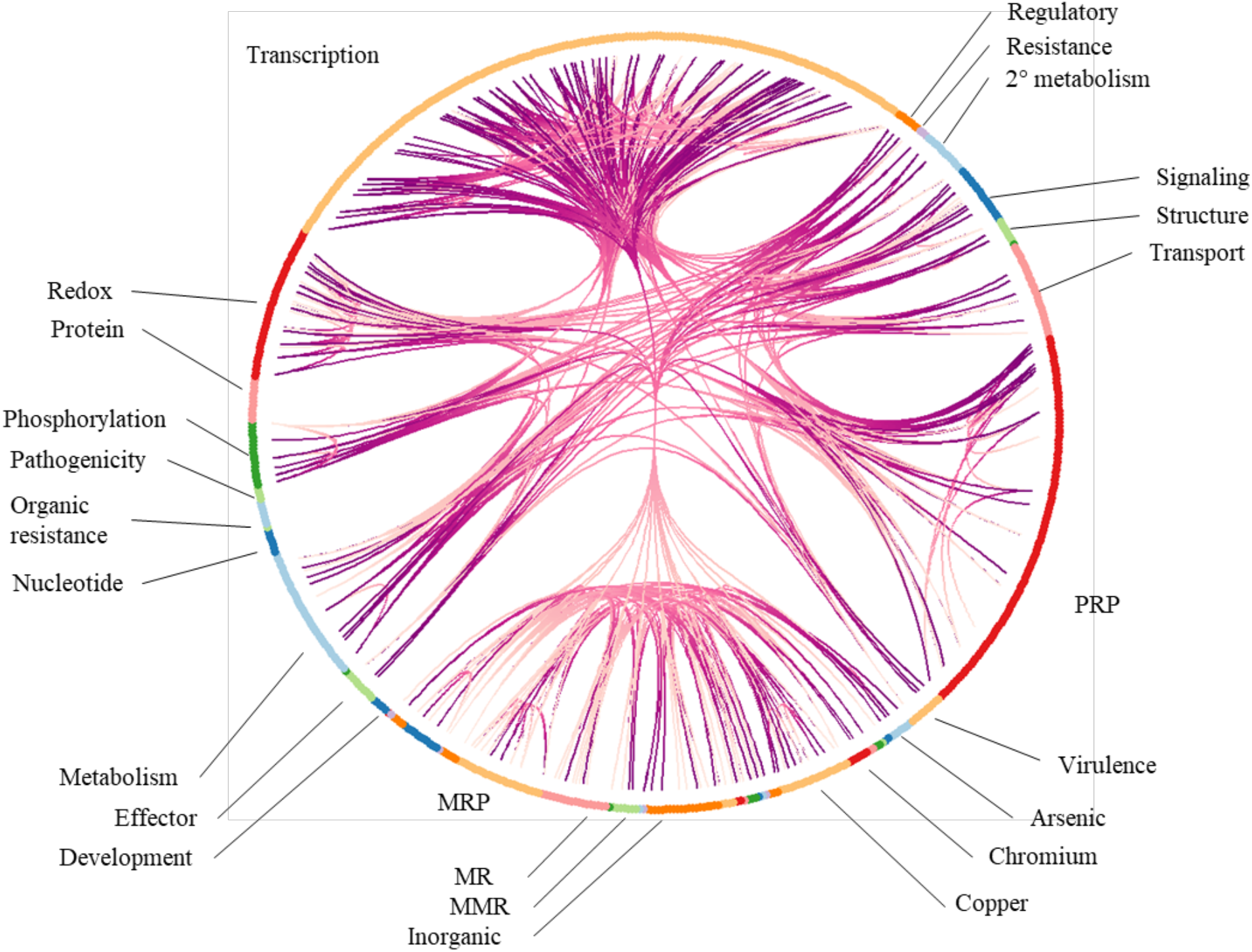
Hierarchical edge network of the different MRPs and PRPs. To avoid overplotting, only the relatively visible PRP and MRP groups are labelled. Each of the nodes at the rim of this diagram represents a PRP or MRP in *Fusarium*. The connections dictate all pairwise cooccurrences that occur at a frequency higher than random chance (p-value < 0.001) as calculated from cooccur package (Veech, 2013).

**Figure 6.**
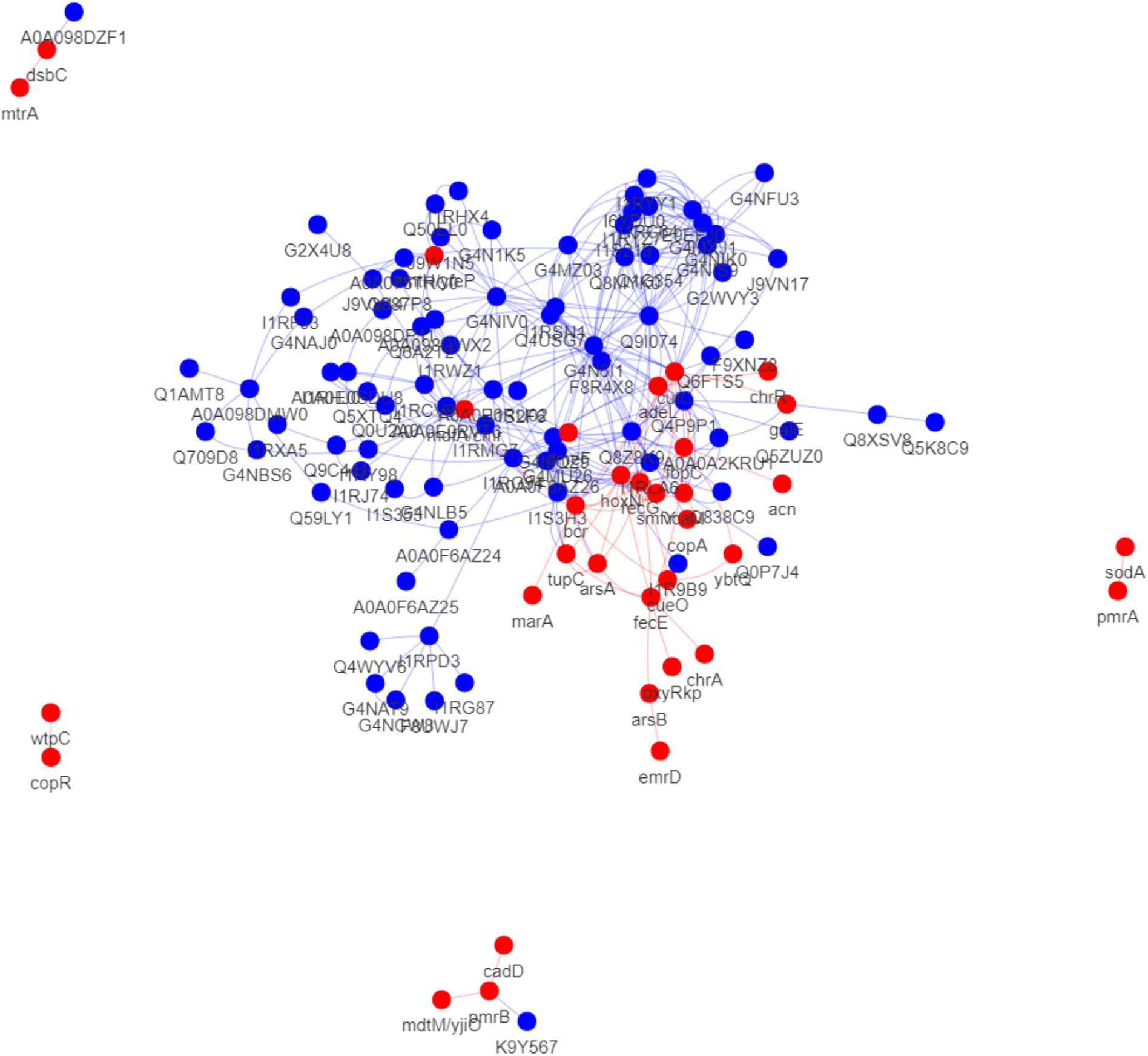
Network diagram. Red nodes represent MRPs while blue nodes represent PRPs. Edges are made along nodes where their observed co-occurrence is greater than chance calculated by cooccur package (Veech, 2013; p < 0.001).

Overall, our results show that there is indeed a significant correlation between MRP and PRP counts. There is also some degree of co-occurrences between both types of proteins. While far from being confirmed, our results suggest further investigation into the possibility that MRP and PRP co-selection or linkage.

There is evidence that exposure to metals increase the likelihood of gene transfer of antibiotic resistance (Zhang et al., 2019; Pal, et al., 2015; Hu, et al., 2016) thus linked sequences can be transferred alongside MRGs. If MRPs and PRPs also co-occur, then analyzing the prevalence of these genes in specific populations can also aid in forming control recommendations to prevent the spread of pathogenic genes, not just antibiotic resistance. Evidence has shown that filamentous fungi have been known acquire pathogenicity determinants via HGT (e.g., Ma, et al., 2010), introgression (e.g., Gladieux et al., 2018), or hybridization (e.g., Feurtey et al., 2019) events. It is possible that MRPs could also be inherited in the same manner and thus, linked transmission with MRPs in this manner could be another avenue for the spread for PRPs in a population. However, direct testing into the transfer of MRPs in fungi is needed.

## Data availability statement

All data analyzed and the supporting information are available in NCBI with accession numbers cited in the manuscript.

## Acknowledgements

The authors would like to thank the following: Accelerated Science and Technology Human Resource Development Program (ASTHRDP) scholarship through the Department of Science and Technology (DOST) that supported the graduate studies of the first author; Drs. Cheng Fang Hong and Ian Kendrich Fontanilla for their valuable insights during the conduct of the study and manuscript writing.

## Supplementary Data

**Table S1.**
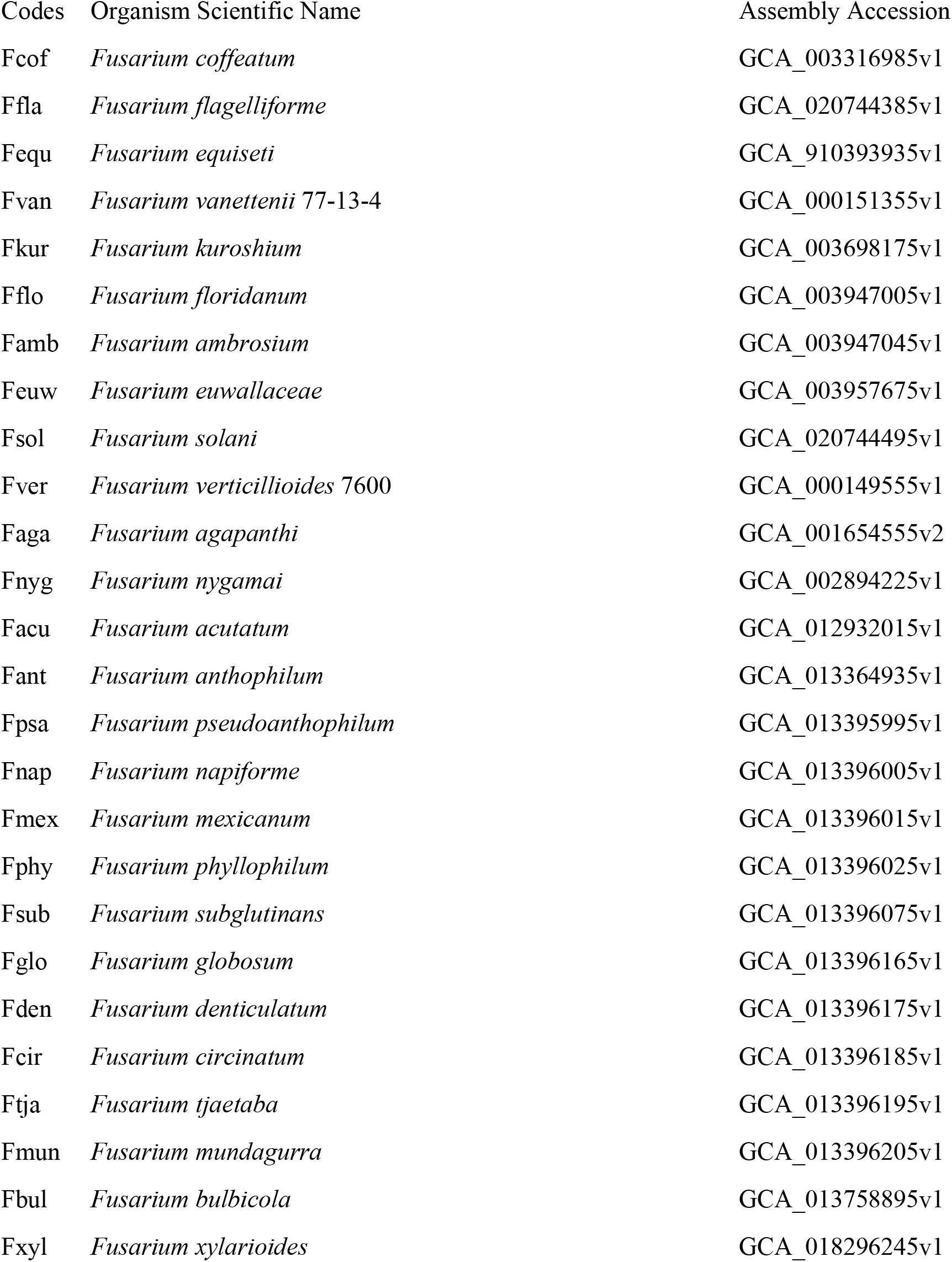
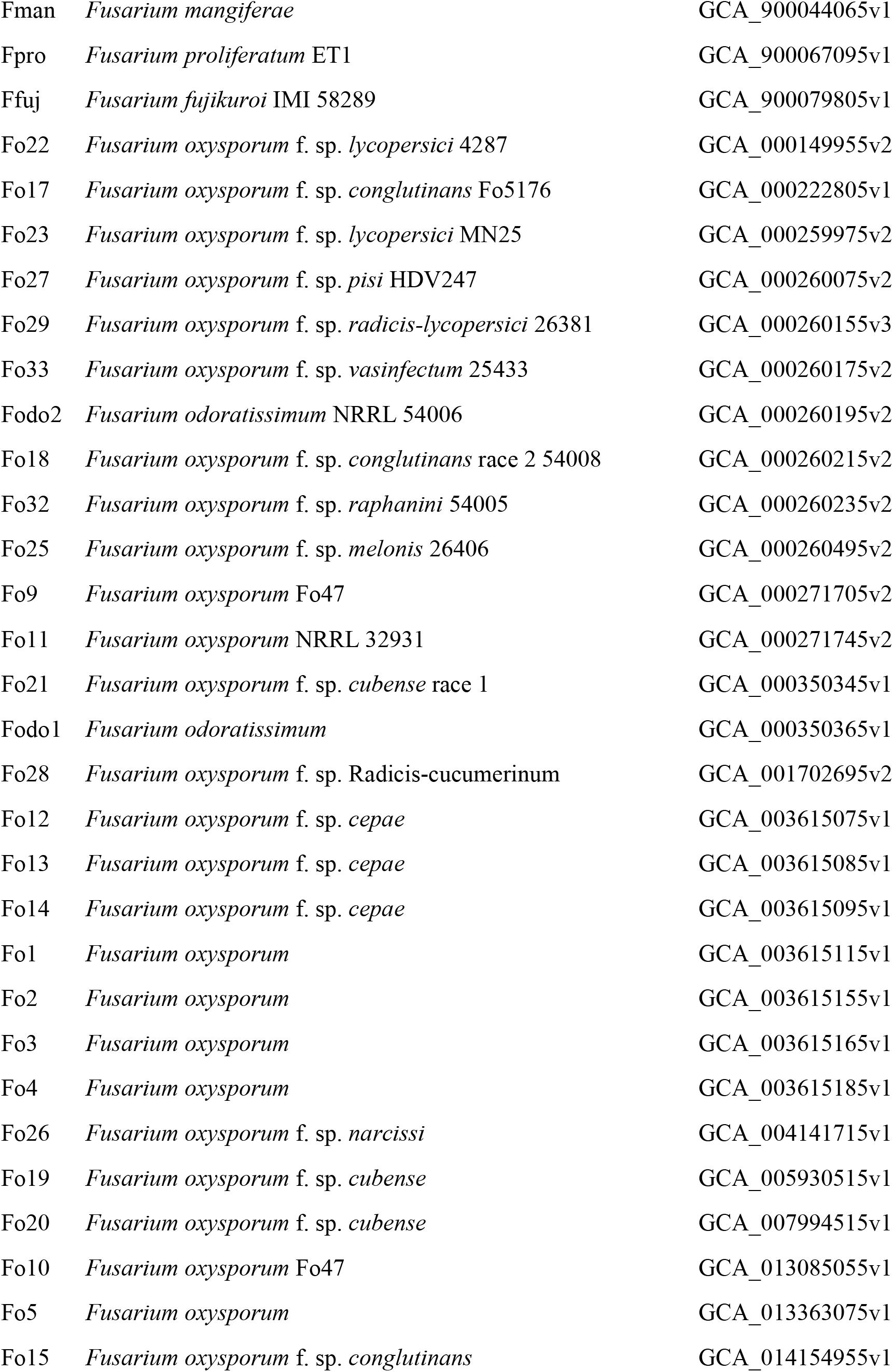
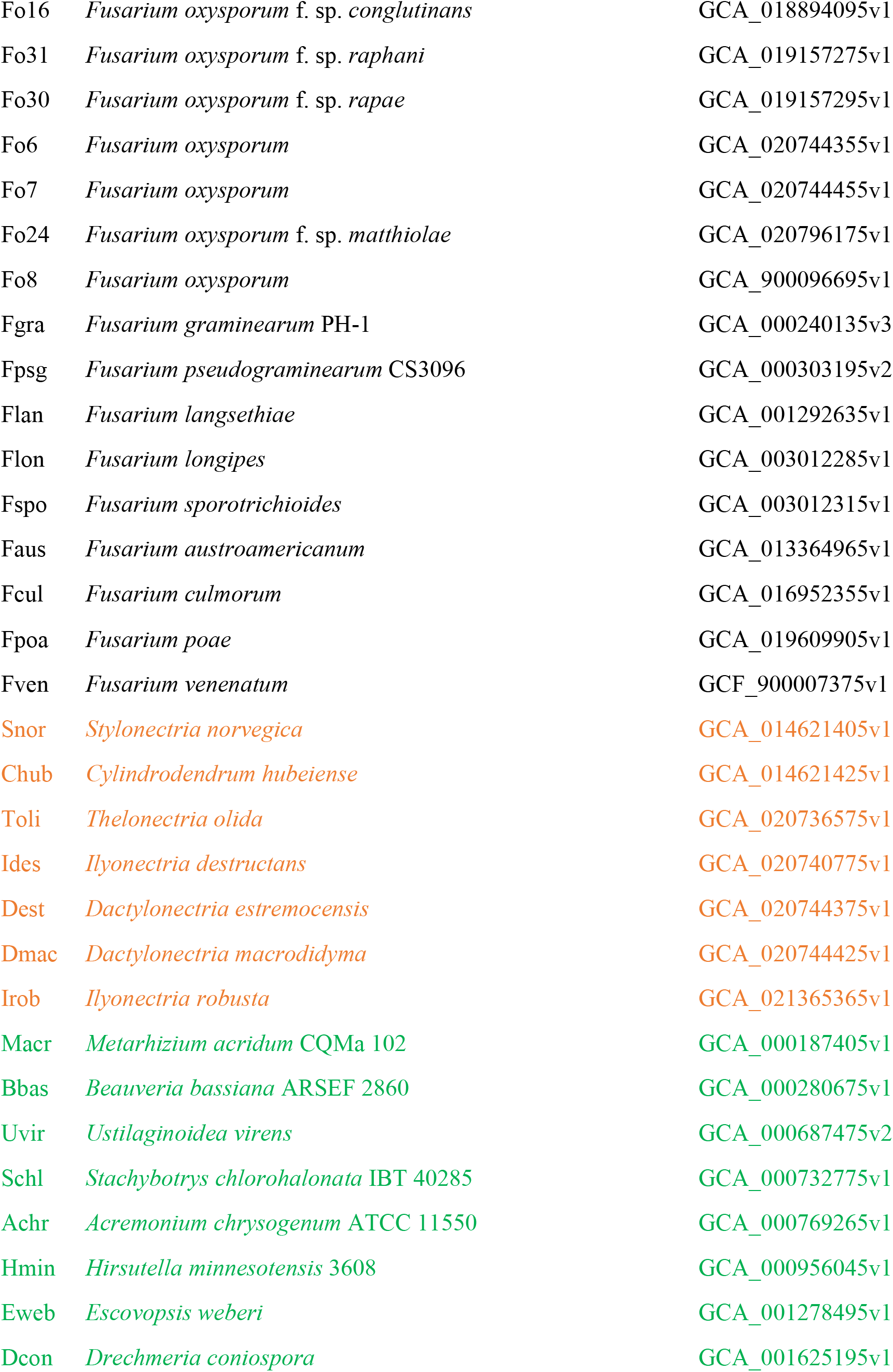
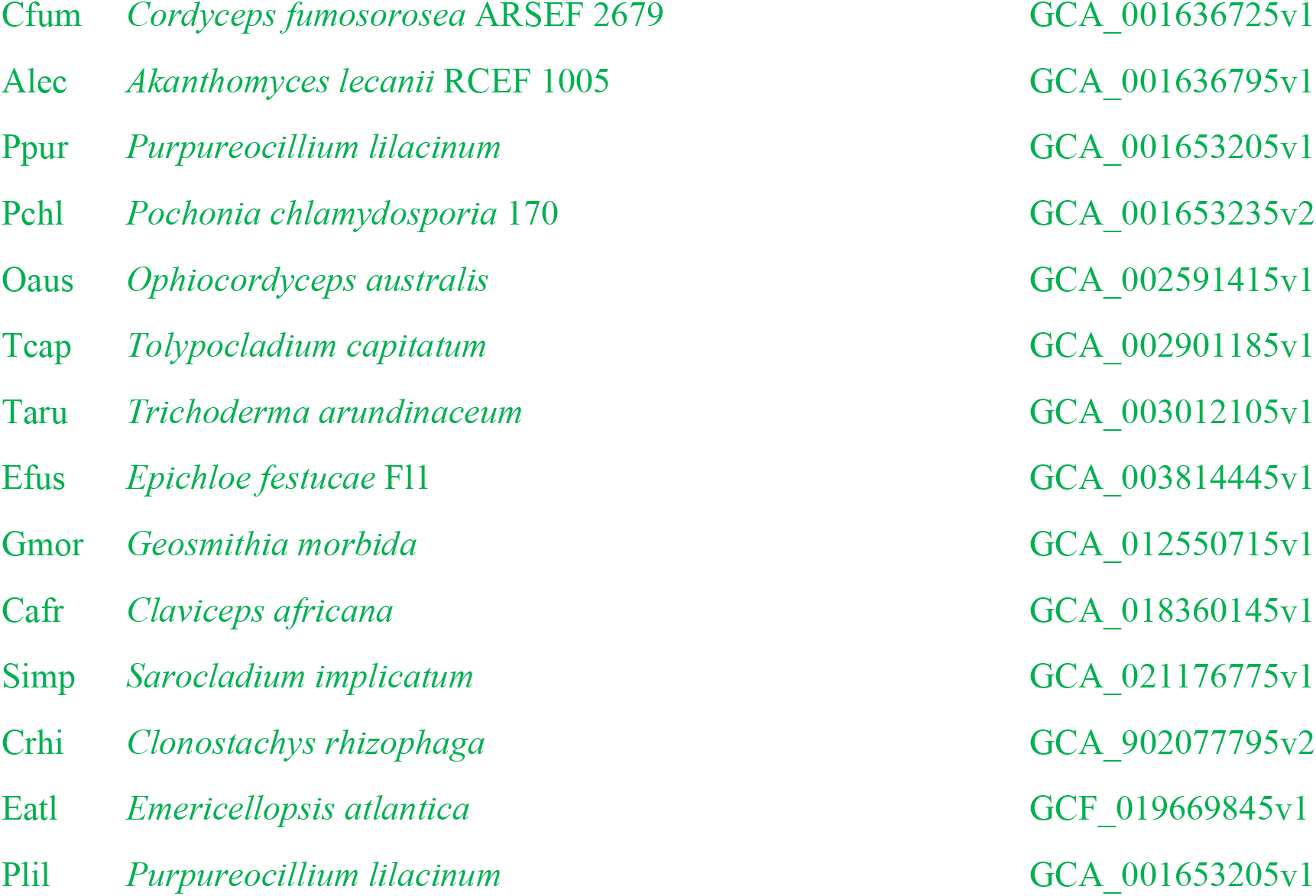
List of organisms referenced in this study (relaxed definition). Taken from NCBI Genome Datasets (https://www.ncbi.nlm.nih.gov/datasets/genomes/). Orange colored names represent members of the family Nectriaceae, while green names represent those included in the Hypocreales.

**Table S2.**
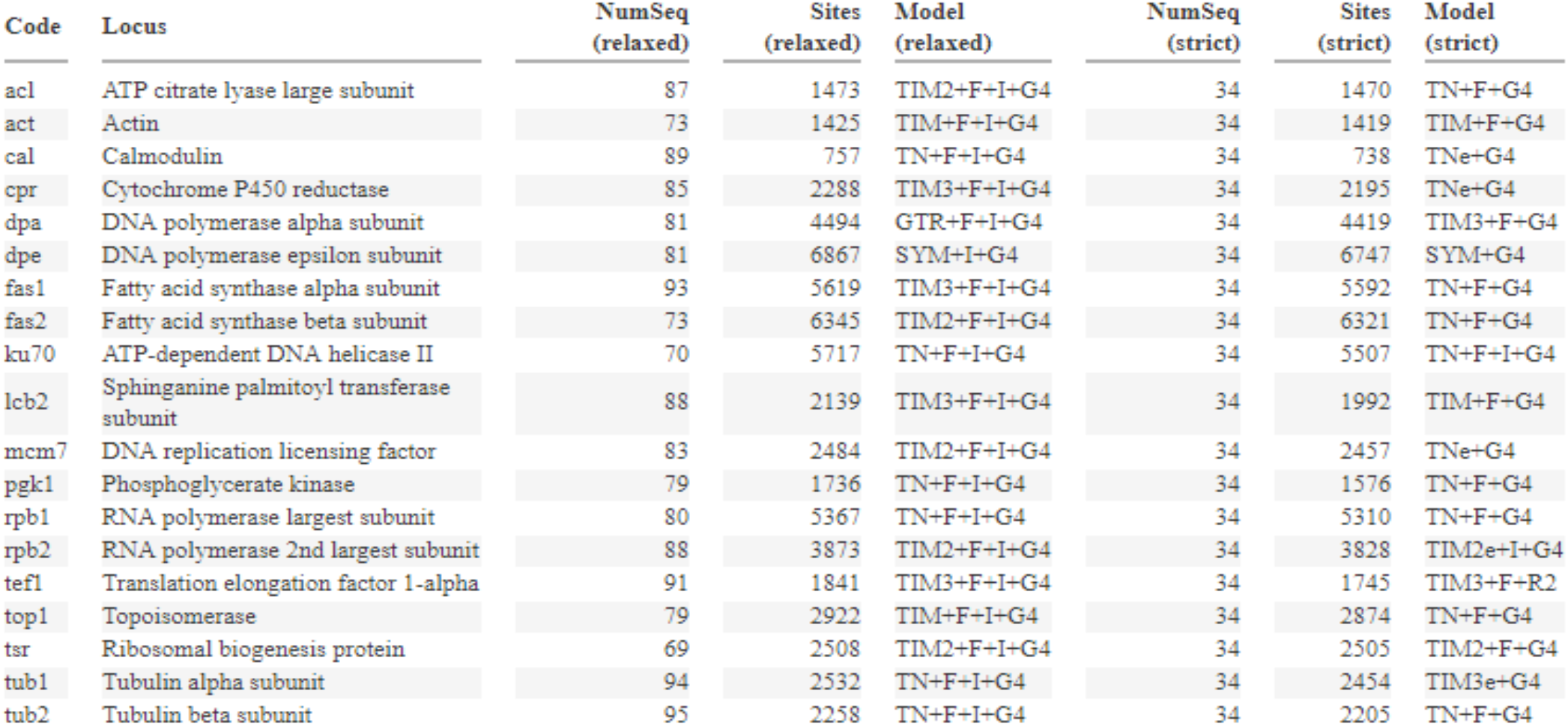
Summary of the 19 loci used for inferring the species tree. Marker choice was adopted from Geiser, et al., (2021). Number of sequences (NumSeq) retrieved from BLASTN hits against the respective target databases and the total number of sites per alignment. Relaxed target database includes all 102 downloaded sequences (see Materials and Methods: Sequence data), strict database includes only species that were not removed due to gaps and saturation (see Strict and relaxed tree definitions below). Table was generated in DAMBE 7 (Xia, 2018). Optimal substitution model was determined using the Bayesian Information Criterion on ModelFinder (Kalyaanamoorthy, et al., 2017). Data were summarized for both relaxed and strict alignments (see below).

**Table S3.**
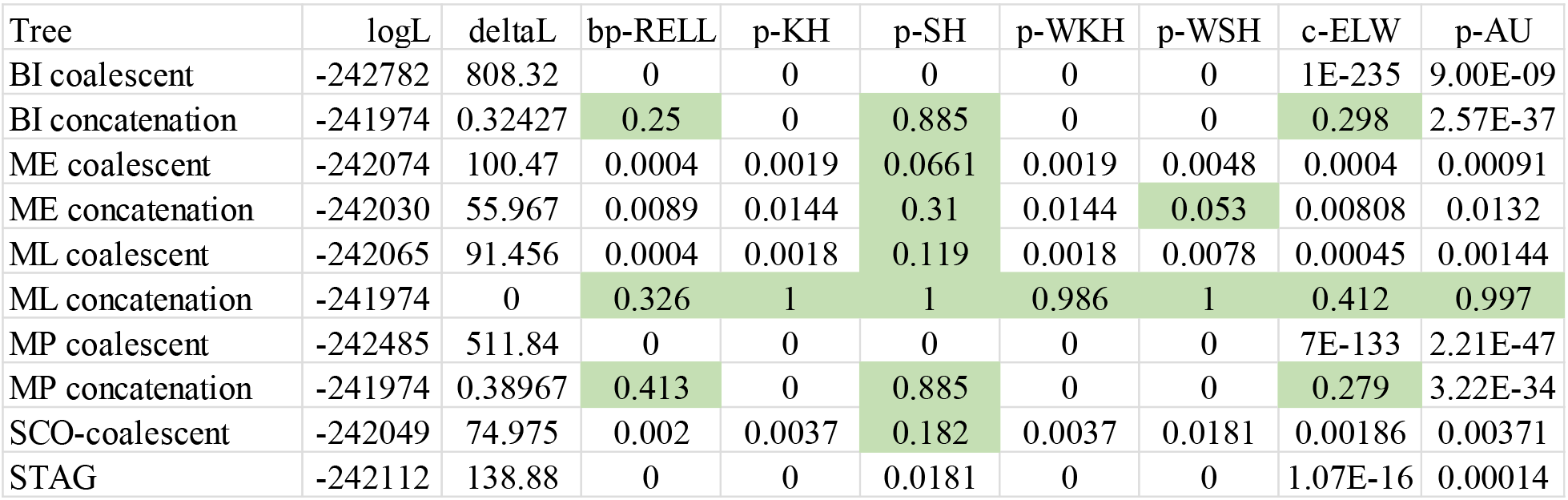
Results of the statistical topology tests performed in IQ-tree2 (Minh, et al., 2020b). Green highlights show that the given tree on the corresponding row cannot be rejected by the null hypothesis (P value > 0.05) – and is therefore equally able to explain the concatenated alignment. Column names are as follows: bp-RELL – RELL bootstrap proportion (Kishino, et al., 1990); p-KH and p-WKH - P -value of unweighted and weighted one sided Kishino-Hasegawa (KH) test (Kishino & Hasegawa, 1989); p-SH and p-WSH—P-value of Shimodaira-Hasegawa test (Shimodaira & Hasegawa, 1999); c-ELW - expected likelihood weight (Strimmer & Rambaut, 2002); p-AU – P – value of approximately unbiased (AU) test (Shimodaira, 2002).

| Tree | logL | deltaL | bp-RELL | p-KH | p-SH | p-WKH | p-WSH | c-ELW | p-AU |
| --- | --- | --- | --- | --- | --- | --- | --- | --- | --- |
| BI coalescent | -242782 | 808.32 | 0 | 0 | 0 | 0 | 0 | 1E-235 | 9.00E-09 |
| BI concatenation | -241974 | 0.32427 | 0.25 | 0 | 0.885 | 0 | 0 | 0.298 | 2.57E-37 |
| ME coalescent | -242074 | 100.47 | 0.0004 | 0.0019 | 0.0661 | 0.0019 | 0.0048 | 0.0004 | 0.00091 |
| ME concatenation | -242030 | 55.967 | 0.0089 | 0.0144 | 0.31 | 0.0144 | 0.053 | 0.00808 | 0.0132 |
| ML coalescent | -242065 | 91.456 | 0.0004 | 0.0018 | 0.119 | 0.0018 | 0.0078 | 0.00045 | 0.00144 |
| ML concatenation | -241974 | 0 | 0.326 | 1 | 1 | 0.986 | 1 | 0.412 | 0.997 |
| MP coalescent | -242485 | 511.84 | 0 | 0 | 0 | 0 | 0 | 7E-133 | 2.21E-47 |
| MP concatenation | -241974 | 0.38967 | 0.413 | 0 | 0.885 | 0 | 0 | 0.279 | 3.22E-34 |
| SCO-coalescent | -242049 | 74.975 | 0.002 | 0.0037 | 0.182 | 0.0037 | 0.0181 | 0.00186 | 0.00371 |
| STAG | -242112 | 138.88 | 0 | 0 | 0.0181 | 0 | 0 | 1.07E-16 | 0.00014 |

**Figure S1.**
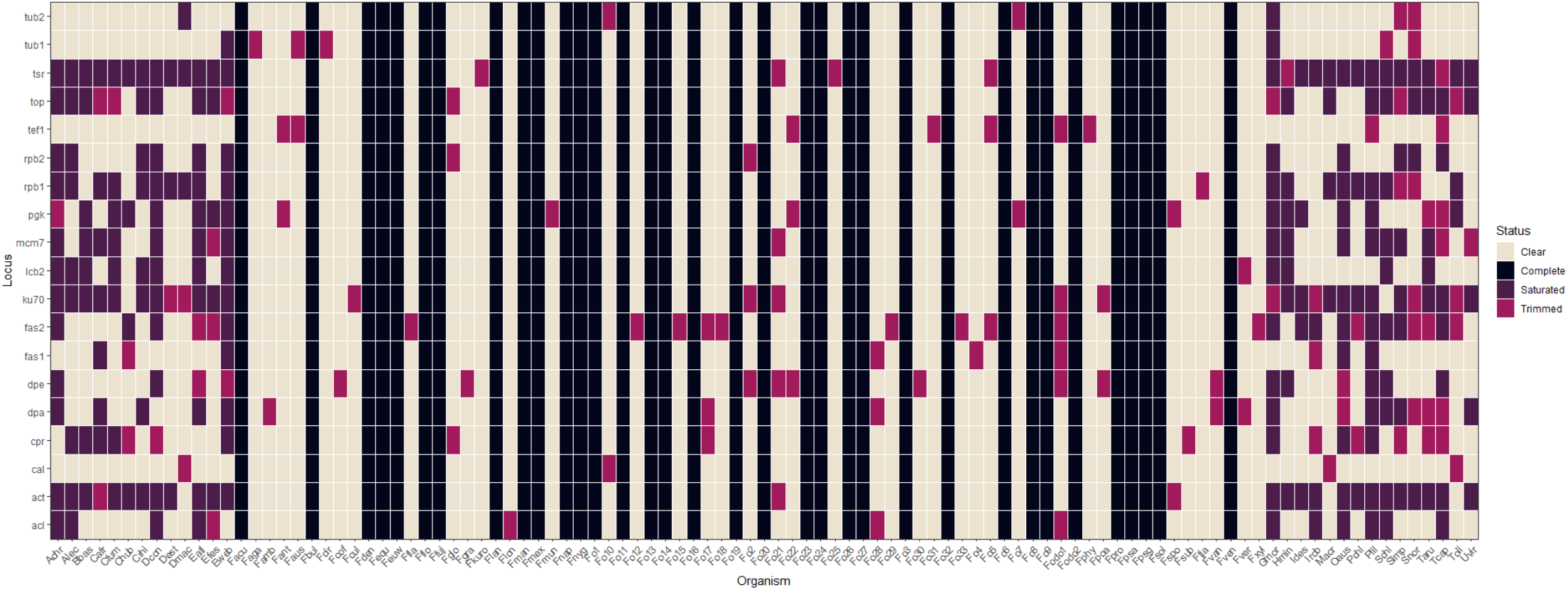
Summary of the loci utilized in the different *Fusarium* species and strains referenced in this paper. All 19 loci were examined for large unique gaps and for substitution saturation (Supplementary Data). Red colors represent loci that were removed due to gaps (trimmed) while purple colors represent loci removed due to substitution saturation (saturated). White (cream) colors represent loci that were not removed. The black vertical colors represent those species with the complete set of loci – and are thus referred to as the ‘strict’ alignments.

**Figure S2.**
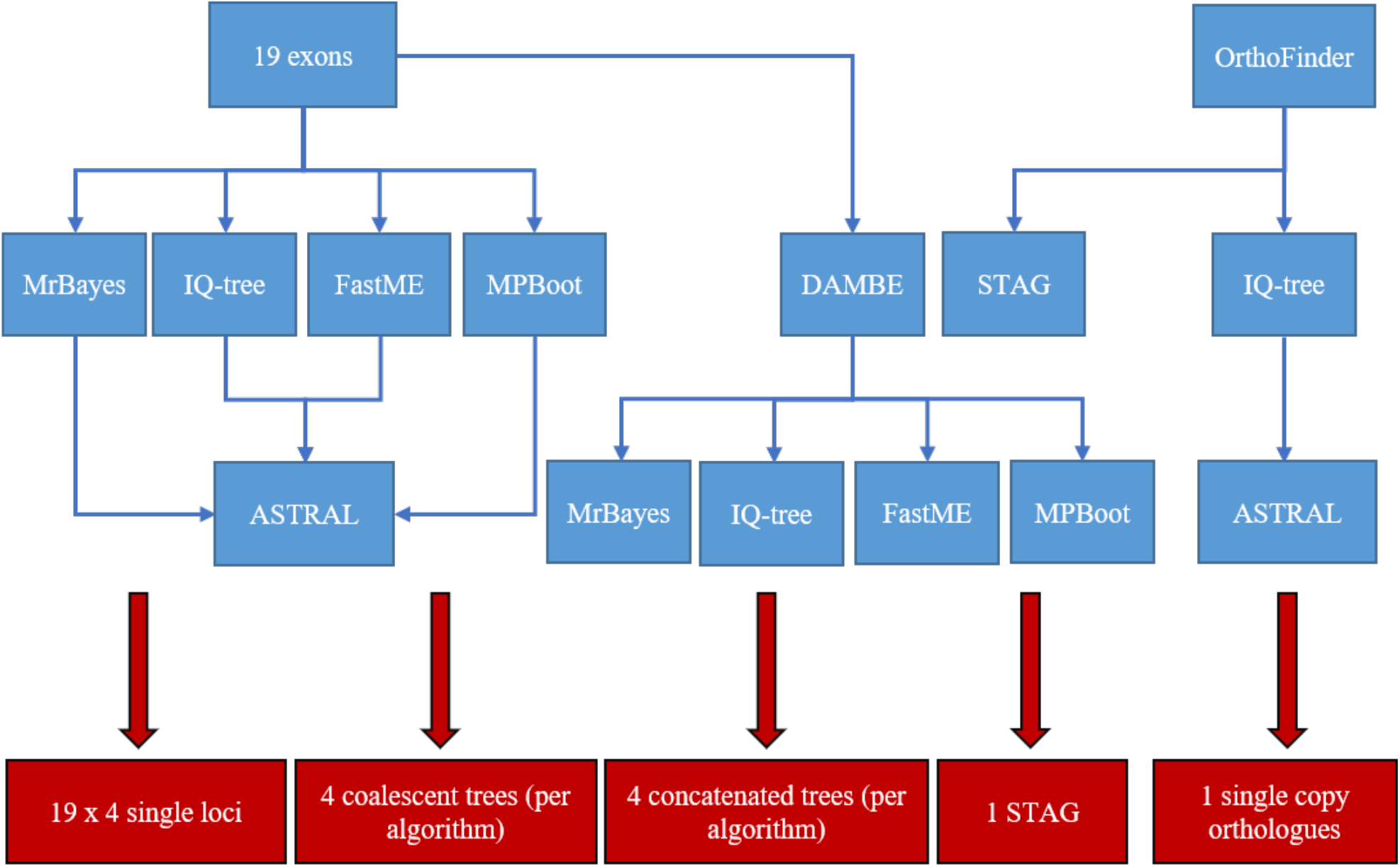
Flowchart of all the phylogenetic trees generated in this study. Red boxes account for the trees used in inferring species phylogeny.

**Figure S3.**
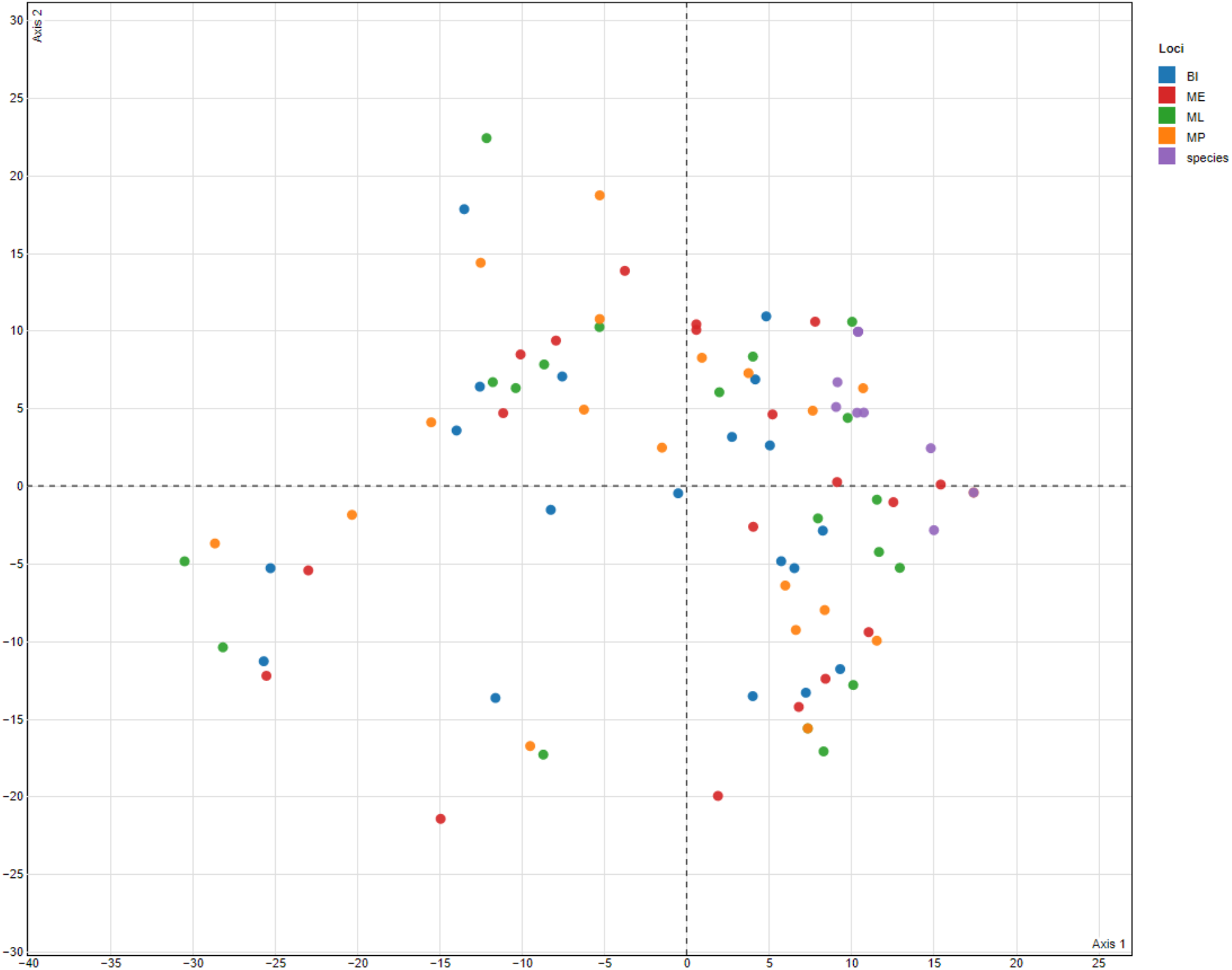
Principal Coordinates Analysis (PCoA) plot of the different trees generated from the 19 loci used for phylogenetic-species identification in Fusarium. Colors correspond to the different clusters formed within the PCA. Clusters were determined using pairwise Euclidean distances within the PCA. Plots were created in treespace (Jombart, et al., 2017). Distances between trees were calculated using the Robinson-Foulds metric (Robinson & Foulds, 1981).

**Figure S4.**
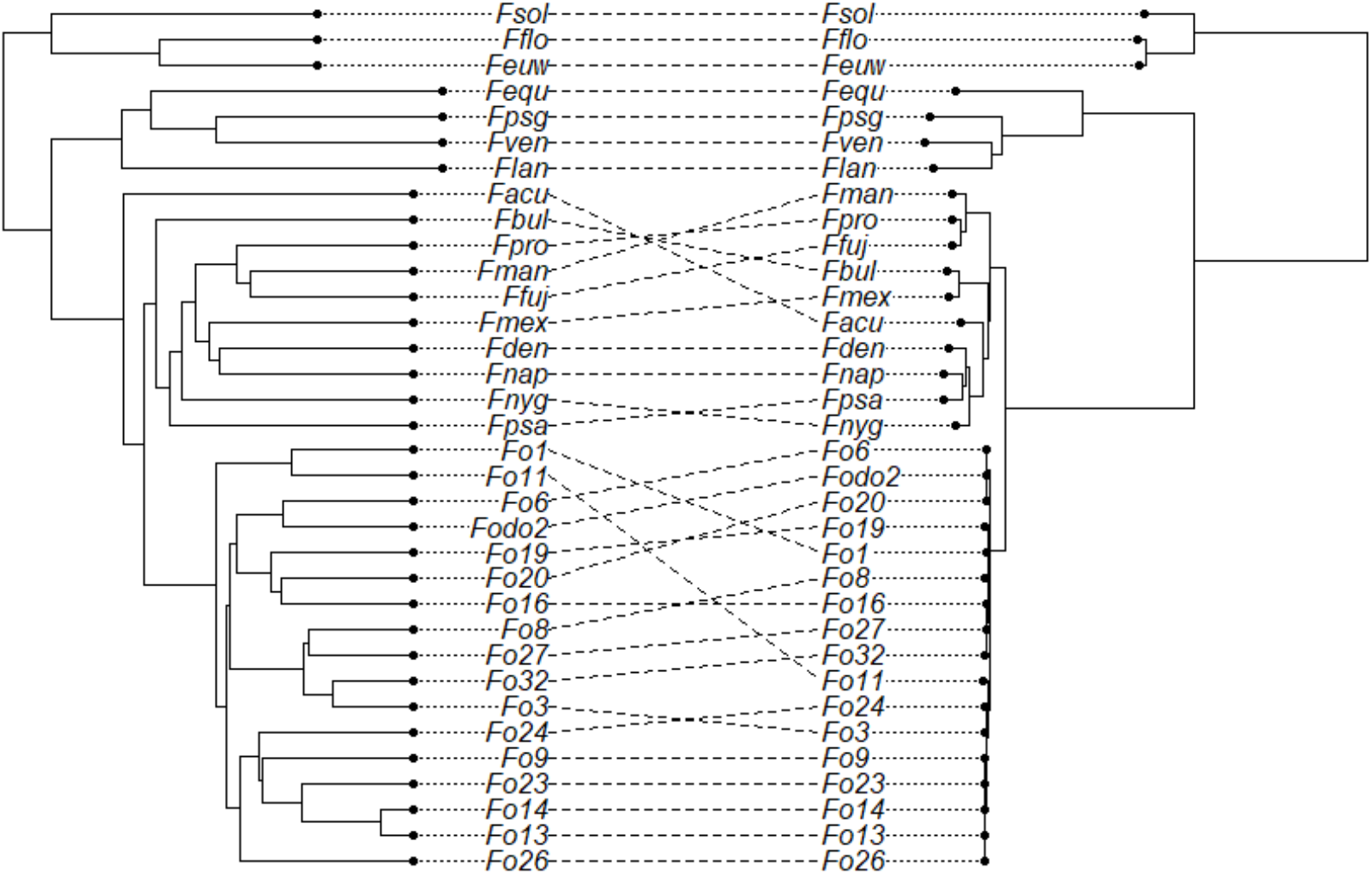
Tanglegram of rooted concatenated-ML tree (right) and MRP genotypes (left) generated using phytools (Revell, 2011). The UPGMA tree is a dendrogram derived from a matrix calculated from the presence/absence of MRPs.

**Figure S5.**
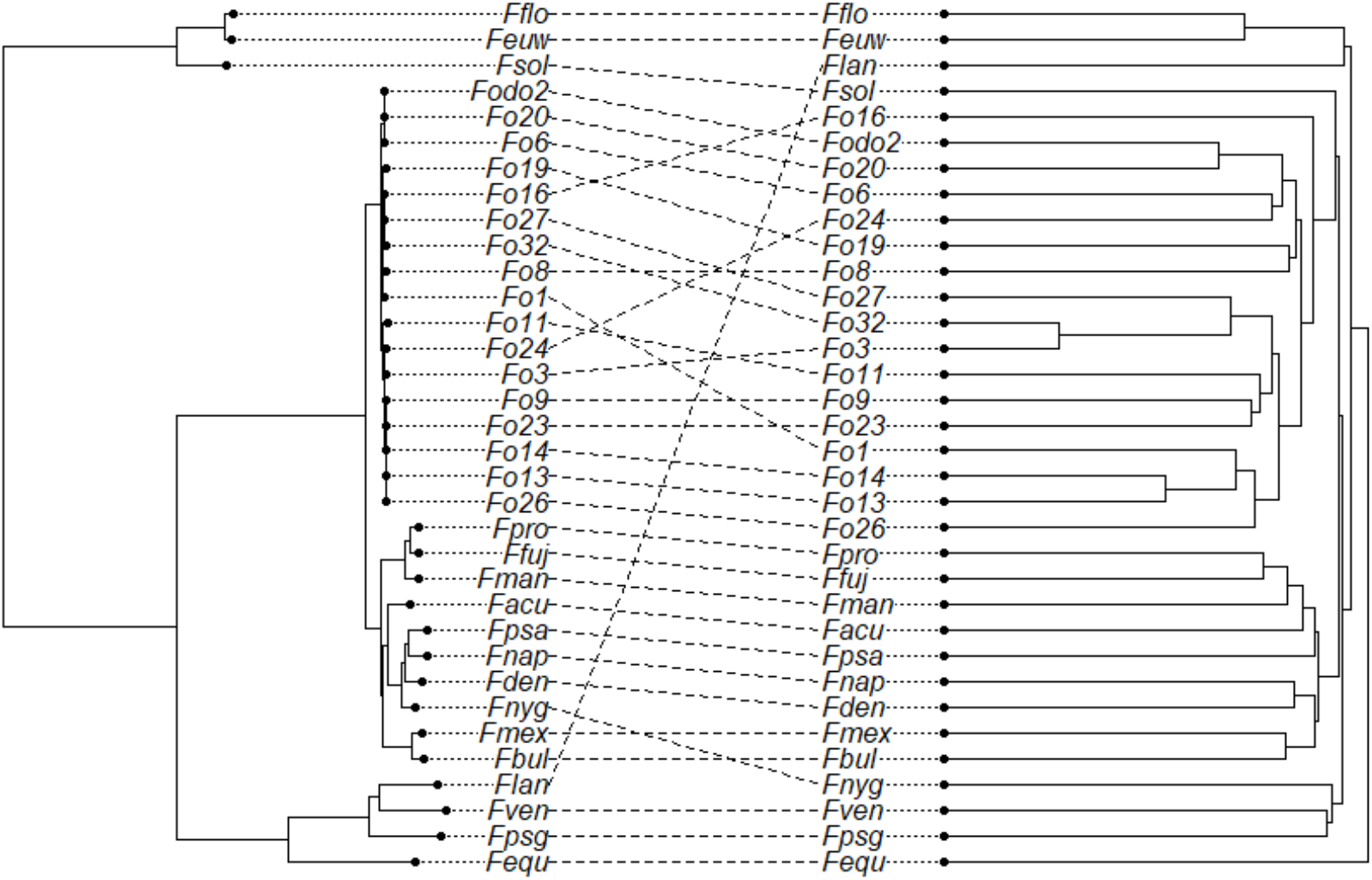
Tanglegram of rooted concatenated-ML tree (left) and PRPs genotypes (right) generated using phytools (Revell, 2011). The UPGMA tree is a dendrogram is derived from a similarity matrix calculated from the presence/absence of PRPs.

**Figure S6.**
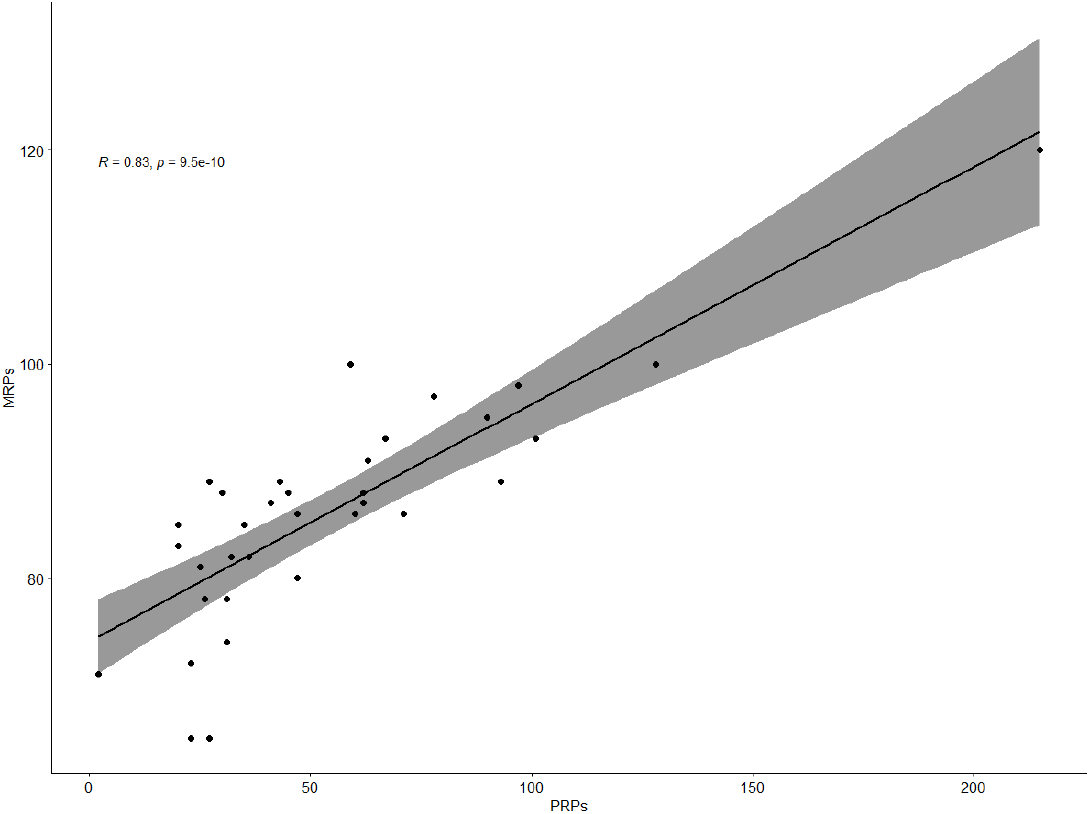
Pearson correlation between the cumulative numbers of MRPs and PRPs per representative proteome. Plot was constructed using R (R Core Team, 2022).

